# Inter-breeder differences in prepulse inhibition deficits of C57BL/6J mice in a maternal infection model for schizophrenia

**DOI:** 10.1101/2020.12.24.423890

**Authors:** Yutaro Kobayashi, Hiroyoshi Inaba, Yuriko Iwakura, Hisaaki Namba, Hidekazu Sotoyama, Yui Murata, Kazuya Iwamoto, Hiroyuki Nawa

## Abstract

Genetic and environmental factors interact with each other to influence the risk of various psychiatric diseases; however, the intensity and nature of their interactions remain to be elucidated. We used a maternal infection model using polyinosinic-polycytidylic acid (Poly(I:C)) to determine the relationship between the maternal breeding environment and behavioral changes in the offspring. We purchased pregnant C57BL/6J mice from three breeders and administered Poly(I:C) (2 mg/kg) intravenously in their tail vein on gestation day 15. The offspring were raised to 8-12 weeks old and subjected to the acoustic startle tests to measure their startle response intensity, prepulse inhibition levels, and degree of the adaptation of the startle response. No statistical interaction between Poly(I:C) administration and sex was observed for prepulse inhibition; thus, male and female mice were analyzed together. The Poly(I:C) challenge significantly decreased prepulse inhibition levels of the offspring born to the pregnant dams from Breeder A but not those from the other breeders. However, there were no significant inter-breeder differences in Poly(I:C) effects on startle response and on startle adaptation. The rearing environment of mouse dams has a prominent impact on the Poly(I:C)-induced prepulse inhibition deficits in this maternal infection model.

## 1. INTRODUCTION

Psychiatric diseases are a major problem for both the affected individual and the society in which they live.^1^ The risk or onset of psychiatric diseases such as schizophrenia is suggested to involve both genetic and environmental factors.^2–4^ Among various environmental factors, maternal infections or inflammation during the gestation draw our attention which can increase the risk of schizophrenia of the child. ^5–7^ Thus, we often employ the rodent offspring whose dams were challenged by viral and bacterial antigens as an animal model for maternal infections or inflammation, which result in various behavioral abnormalities relevant to schizophrenia endophenotypes. ^8–11^ Such associations warrant a discussion on the biological links between the maternal immune inflammatory history and the increased risk of the offspring developing this disease. Previous neuropathological studies on this maternal infection model have reported that the brains of the neonates born to the mothers exhibit the increased cell density or pathological abnormalities of the cerebral cortex. ^12,13^ Poly(I:C) is the double-stranded RNA analog for viral genome, which is recognized by toll-like receptor 3 to trigger the immune inflammatory responses to a viral infection.^14^ Biological factors that influence the neurobehavioral consequences of the Poly (I:C) challenge have been reported from various perspectives.^15–18^ The rearing environment of mice in the laboratory alters the neurobehavioral effect of Poly (I:C).^19^ The environment also has marked impact on the behavioral traits and immune inflammatory responses of mice themselves.^20,21^ In this study, we purchased pregnant mice with the same C57Bl/6J genetic background from three Japanese breeders and treated them with Poly(I:C) to investigate how environmental factors (i.e., breeder) affected the behavioral traits of their offspring at maturity.

## 2. MATERIALS AND METHODS

### 2.1 Animals

The genetic strain of C57Bl/6J mice was used in this experiment. Six to ten pregnant mice were purchased from CLEA Japan Inc. (Tokyo, Japan) and grown in their Fuji breeding factory, from Charles River Laboratories Japan Inc. (Kanagawa, Japan) and grown in their Hino breeding factory, and from SLC Japan Inc. (Shizuoka, Japan) and grown in their Haruno breeding factory. In the authors’ institute, the dams and their pups were kept in a specific-pathogen-free environment, and the room temperature and humidity were maintained at 23℃ and 50–70%, respectively, with a constant 12-h light-dark cycle (lights off: 8:00 PM-8:00 AM). All animals were provided the same feed (NMF, Oriental Yeast, Inc., Tokyo, Japan) and ad libitum access to water. All experiments adopted in this study were approved by and conducted under the guidance of the Niigata University Animal Ethics Committee.

### 2.2 Poly(I:C) administration

Pregnant mice on gestation day 15 underwent intravenous injection with 2.0 mg/kg Poly(I:C) (potassium salt, G.E. Healthcare, Illinois, USA) or vehicle (sterile pyrogen-free 0.9% NaCl).^22,23^ To obtain the final Poly(I:C) concentration of 1 mg/mL, it was dissolved in sterilized 0.9% NaCl without pyrogenic substances. The Poly(I:C) solution or saline was administered to the tail vein under mild physical restraint. The pups were weaned on postnatal day 21, and 2–3 of each sex was randomly assigned to each group. The number of total offspring born was 159 pups.

### 2.3 Prepulse inhibition test

The pups’ acoustic startle response, startle adaptation, and prepulse inhibition (PPI) were assessed between postnatal days 56 and 84.^24^ Mice were placed in a plastic cylinder, and the cylinder was fixed to a testing chamber (SR-Lab Systems, San Diego, CA, USA). After completing the acclimatization period with 70 dB background noise (white noise), the main acoustic stimuli (120 dB, white noise; duration, 40 ms) were presented with prepulse stimuli (73, 76, 79, or 82 dB) to mice in an order predetermined by a pseudorandom number generator with 15-s intervals between stimulus presentations.^24^ PPI was measured conforming to a previous report.^24^ Before measuring PPI, a 120-dB acoustic stimulus was administered to mice five times and the proportion of the startle change, was calculated as an adaptation (%).

### 2.3 Sound startle response test

Mice were placed in a plastic cylinder which was fixed to a testing chamber (SR-Lab Systems, San Diego) as described above. Acoustic stimuli (90, 95, 100, 105, 110, 115, and 120 dB, white noise; duration, 40 ms) were presented in a pseudorandom manner with 15-s intervals.^24^ Their mean intensity for 120-dB stimuli were selected and presented as the intensity of the startle response.

### 2.4 Statistical analysis

Factorial analysis of variance (ANOVA) was first performed with the main factors of Poly (I:C), breeder, tone intensity, and sex using SPSS ver. 11.0 (SPSS Japan, Tokyo, Japan). As there was a significant interaction between breeder and other indices, following statistical analyses were done in each breeder. Interactions between behavioral data and sex were also analyzed (Supplementary Table 1). In the absence of such interaction, the data from both sexes were combined. Post-hoc analyses were performed by planned comparisons of Welch t-test with Holm’s correction or by multiple comparisons of Tukey–HSD test. *P*< 0.05 was considered statistically significant.

## 3. RESULTS

### 3.1 Prepulse inhibition difference among mouse breeders

PPI levels are known to decrease in patients with various psychiatric diseases such as schizophrenia and thus we employed it as behavioral markers for its animal modeling. ^24–26^ The effects of the maternal Poly(I:C) challenge on PPI was measured when pups reached maturity (Figure 1). First, we performed factorial ANOVA with the between subject factors of breeders, Poly(I:C), and sex, and the within-subject factors of prepulse. We found a significant interaction between breeders and Poly(I:C) (*F_2,130_* = 3.206, *P* = 0.044) and no significant interaction between sex and any other subject factor, suggesting that Poly(I:C) effects varied among breeders but not between the sexes. Thus, we divided the statistical analyses into individual breeders, combining the male and female PPI data (see statistical details in Supplemental Table 1). PPI levels only decreased significantly in the pups of C57BL/6J dams purchased from CLEA Japan (*F_1,52_* = 5.659, *P* = 0.021). When pregnant C57BL/6J dams purchased from Charles River Japan and SLC Japan, the PPI of their pups was not significantly altered by Poly(I:C) treatment (Charles River: *F_1,31_* = 0.725, *P* = 0.401; SLC: *F_1,47_* = 1.960, *P* = 0.168).

**Figure 1.**
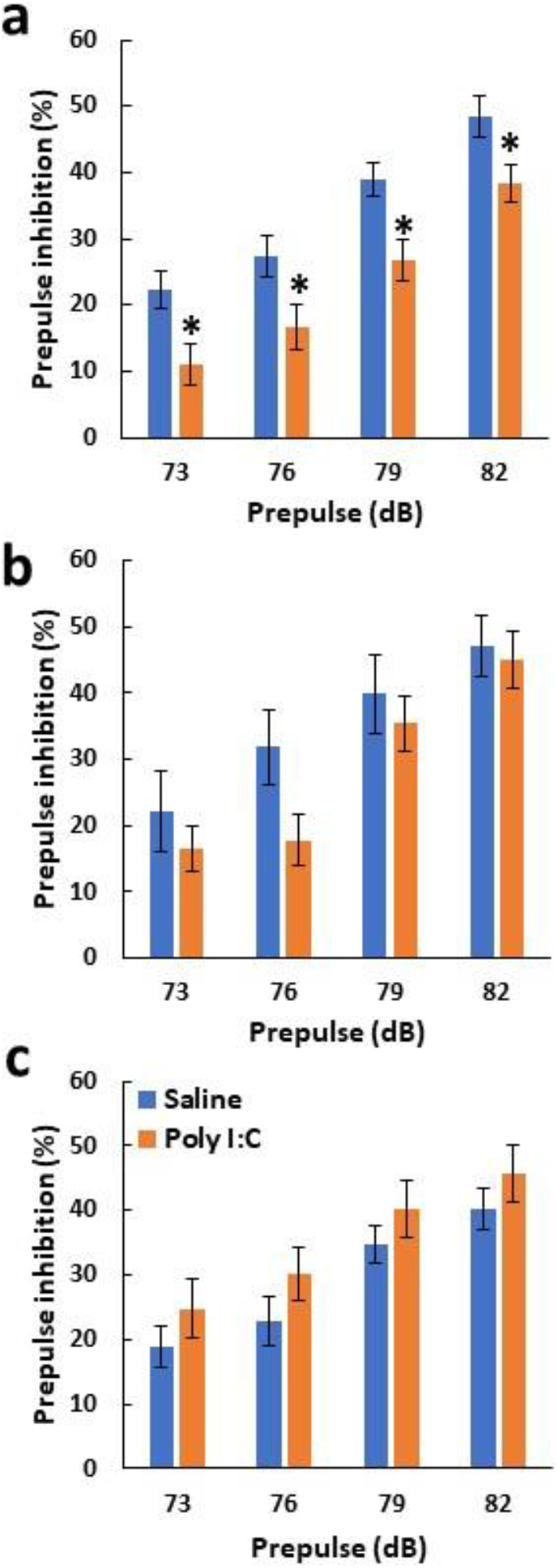
The effect of breeders on prepulse inhibition after administering Poly(I:C) to their dams. The inhibitory effects of prepulse sounds at 73, 76, 79 and 82 dB were assessed as PPI levels in pups born to pregnant C57BL/6J mice after raising them for 2–3 months. Pregnant mice were purchased from CLEA Japan (a), Charles River Japan (b), and SLC Japan (c) and challenged with Poly(I:C) or saline on gestation day 15 (n = 3–6 dams for each group). Data are presented as mean ± SEM. PPI was observed in the inhibitory % of the main pulse (120 dB) (14–39 pups for each group). \**P* < 0.05, planned multiple comparison was performed between saline and Poly(I:C) groups at each prepulse levels using Welch t-test with the Holm’s compensation.

### 3.2 Influences of mouse breeders on sound startle response and test adaptation

At maturity, the Poly(I:C) effect on the offspring startle response was also measured with 120 dB sound. Factorial ANOVA was similarly performed with the between-subject factors of breeders, Poly(I:C), and sex. There were main effects of breeder (*F_2,130_* = 5.68, *P* = 0.004) and sex (*F_1,130_* = 20.2, *P* < 0.001) but not that of Poly(I:C) (*F*1,130 = 0.892, *P* = 0.347) with no interactions between any subject factors (Supplemental Table 1). Accordingly, the intensity of sound startle responses was plotted in each breeder and each sex (Figure 2a-c). Post-hoc analyses detected the significant sex difference in all breeders (CLEA: *P* = 0.012, Charles: *P* = 0.028, SLC: *P* = 0.02) and found that the startle response of the offspring from CLEA Japan was significantly higher than that from SLC Japan (*P* = 0.002). Although there were differences in basal startle amplitudes among breeders and sex, the behavioral scores were not affected by administering Poly(I:C) to the dam during pregnancy.

**Figure 2.**
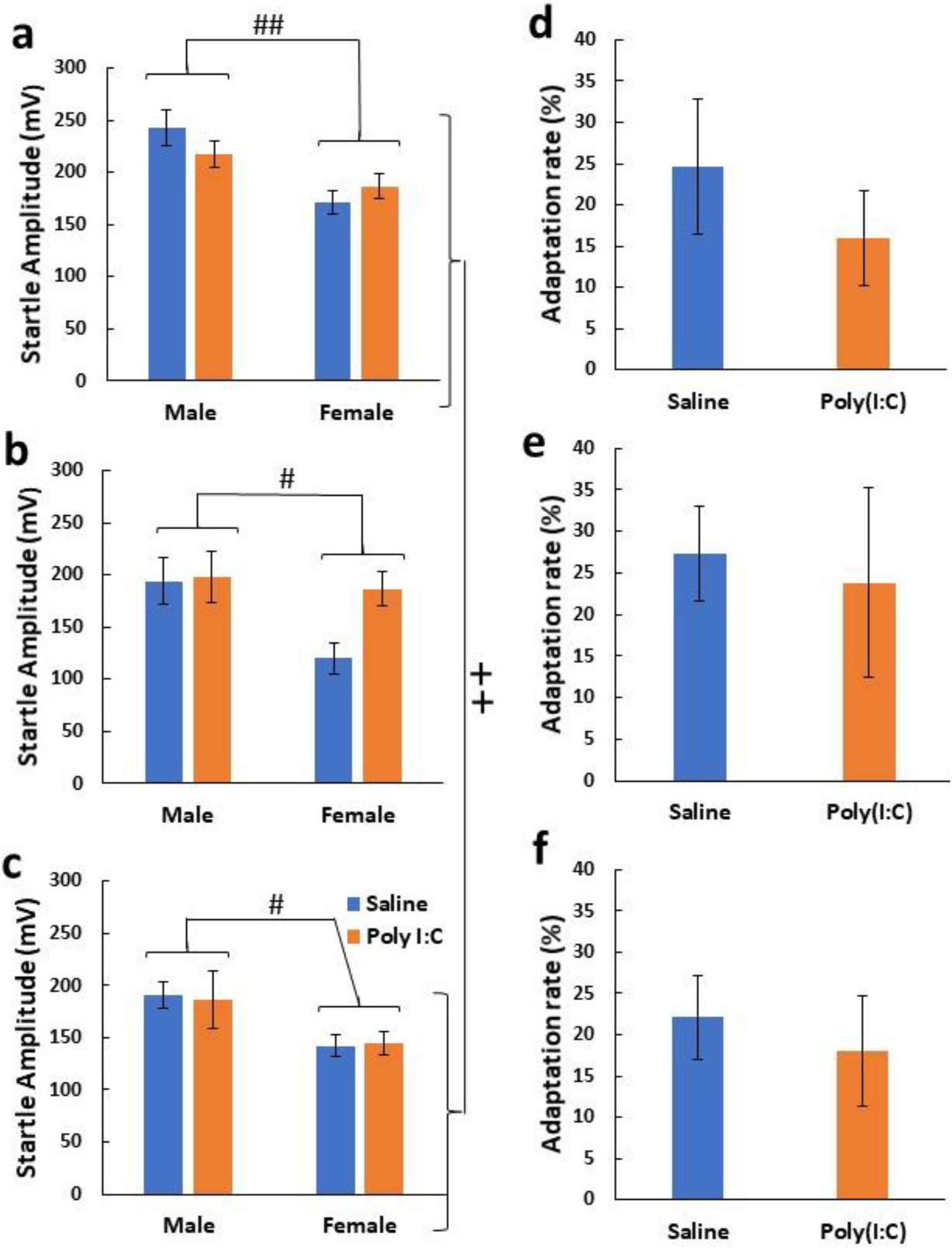
Breeder effects on sound startle response and adaptation. Employing the same sets of mice in Figure 1, we determined the intensity of their startle response (mV) to 120-dB white noises (a–c) and adaptation rates of the startle responses (%) to the 120-dB noises (d–f). The pregnant mice (C57BL/6J) were purchased from CLEA Japan (a, d), Charles River Japan (b, e), and SLC Japan (c, f). Data are presented as mean ± SEM. See statistical details in Supplementary Table 1. In the startle intensity, there were significant an inter-breeder difference between CLEA Japan and SLC Japan (^++^*P* < 0.01, Tukey–HSD) and a sex difference in all breeders (^#^*P* < 0.05, Welch t-test with the Holm’s compensation), which was irrespective of the Poly(I:C) effects.

It has been reported that the ability to adapt to external stimuli is reduced in patients with psychiatric disorders.^27,28^ The effects of Poly(I:C) on the adaptation of pups for PPI test paradigm were assessed at their maturity (Figure 2d-f). Factorial ANOVA found no significant effects of breeders, Poly(I:C), or sex with no their interactions (Supplemental Table 1).

## 4. DISCUSSION

In the present study, we focused on one of the suggested environmental factors, maternal breeding environment, to investigate how this environmental factor affects the sound startle feature of this, model; PPI levels, sound startle responses, and their startle adaptation.^8,22,23,29^ We purchased dam mice with the same genetic background that were raised in three different environments and observed the following behavioral similarity and difference of their offspring across dam’s breeders: (1) Maternal Poly (I:C) challenge significantly disrupted PPI levels of their offspring only when pregnant mice were purchased from CLEA Japan; (2) The Poly (I:C) effects on the startle responses of their offspring were indistinguishable among all companies, and (3) The adaptation to acoustic stimuli did not differ among any origins of the companies, (4) Irrespective of the Poly (I:C) effects, the basal behavioral feature of sound startle response varied among breeders and between the sexes.

Challenging mouse dams with Poly(I:C) had different effects on pups’ PPI levels depending on the breeder that the dam was purchased from. This presumably represents a result of the biological interaction between genetic factors and breeding environment (i.e., environmental factors) of the pregnant mice. This finding appears to agree with the report that the cognitive-behavioral changes such as sociability and exploratory behavior in pups induced by Poly(I:C) injection varied between the breeders that provided the dams.^30^ The present inter-breeder difference is consistent with the fact that the blood levels of Poly(I:C)-induced inflammatory cytokines differ depending on the dam’s breeding environment.^20,30^

In this experiment, Poly(I:C) was administered to pregnant mouse dams (the C57BL/6J strain) at the mid-to-late pregnancy period and exerted the PPI effects on their offspring. This may be in agreement with the previous study that PPI levels were decreased in the pups of pregnant mice (the genetic strain of C57BL/6J) challenged intraperitoneally with Poly(I:C) on the late gestation.^22^ However, another study reported no changes in PPI in the pups of pregnant mice (C57BL/6J) when treated with the higher dose of Poly(I:C) on the late gestation stage.^15^ This controversy suggests the possibility that the present breeder effects could be influenced by the experimental conditions such as Poly(I:C) doses, its injection sites, pregnancy periods, etc.

In summary, this study revealed that the breeder from which mice are procured is an important factor in the mouse model for behavioral deficits produced by Poly(I:C). Further investigations on the specific aspects of the breeding environment underlying our findings are reflected in mice *in vivo* are warranted.

## ACKNOWLEDGMENTS

This study was supported by MEXT Grant-in-Aid for Scientific Research on Innovative Areas “Multiscale Brain” (HNawa: 18H05429, 18H05428, KI: 18H05430), AMED (KI: JP20dm0207074) and by The Uehara Memorial Foundation to HN.

## AUTHOR CONTRIBUTIONS

YK performed the experiments and coordinated the work presented. HNawa designed the experiments and wrote the manuscript. HNamba, YI, HN, HS, YM, and KI provided technical assistances and commented on the manuscript

