## Supplemental Table 1 for "Inter-breeder differences in prepulse inhibition deficits of C57BL/6J mice in a maternal infection model for schizophrenia"

**Supplementary Table S1**; the sample number of mouse dams and pups born

Main effects and interactions of the subject factors of breeder (3), Poly(I:C)(2), and sex (2)

| Subject | Breeder | The number of the dam/ The number of the offspring | Factors | F value and p value |
|---|---|---|---|---|
| Prepulse inhibition | All | Saline : 11/♂ : 30, ♀ : 35<br>Poly(I:C) : 14/♂ : 40, ♀ : 37 | Breeder | F(2, 130)=0.760, p=0.470 |
| | | | Poly(I:C) | F(1, 130)=1.236, p=0.268 |
| | | | Sex | F(1, 130)=0.098, p=0.754 |
| | | | Interaction(Breeder X Poly(I:C)) | F(2, 130)=3.206, p=0.044 |
| | | | Interaction(Breeder X Sex) | F(2, 130)=0.762, p=0.469 |
| | | | Interaction(Breeder X Poly(I:C)) | F(1, 130)=0.014, p=0.906 |
| | | | Interaction<br>(Breeder X Poly(I:C) X Sex) | F(2, 130)=0.132, p=0.876 |
| | CLEA | Saline : 3/♂ : 7, ♀ : 10<br>Poly(I:C) : 6/♂ : 23, ♀ : 16 | Poly(I:C) | F(1, 52)=5.659, p=0.021 |
| | | | Sex | F(1, 52)=0.005, p=0.944 |
| | | | Interaction(Poly(I:C) X Sex) | F(1, 52)=0.164, p=0.687 |
| | Charles | Saline : 3/♂ : 11, ♀ : 10<br>Poly(I:C) : 3/♂ : 8, ♀ : 6 | Poly(I:C) | F(1, 31)=0.725, p=0.401 |
| | | | Sex | F(1, 31)=0.155, p=0.697 |
| | | | Interaction(Poly(I:C) X Sex) | F(1, 31)=0.075, p=0.786 |
| | SLC | Saline : 5/♂ : 12, ♀ : 15<br>Poly(I:C) : 5/♂ : 9, ♀ : 15 | Poly(I:C) | F(1, 47)=1.960, p=0.168 |
| | | | Sex | F(1, 47)=1.654, p=0.205 |
| | | | Interaction(Poly(I:C) X Sex) | F(1, 47)=0.071, p=0.792 |
| Startle respnse | All | Saline : 11/♂ : 30, ♀ : 35<br>Poly(I:C) : 14/♂ : 40, ♀ : 37 | Breeder | F(2, 130)=5.680, p=0.004 |
| | | | Poly(I:C) | F(1, 130)=0.892, p=0.347 |
| | | | Sex | F(1, 130)=20.188, p<0.001 |
| | | | Interaction(Breeder X Poly(I:C)) | F(2, 130)=1.383, p=0.254 |
| | | | Interaction(Breeder X Sex) | F(2, 130)=0.054, p=0.947 |
| | | | Interaction(Poly(I:C) X Sex) | F(1, 130)=3.157, p=0.078 |
| | | | Interaction<br>(Breeder X Poly(I:C) X Sex) | F(2, 130)=0.613, p=0.543 |
| Adaptation rate | All | Saline : 11/♂ : 30, ♀ : 35<br>Poly(I:C) : 14/♂ : 40, ♀ : 37 | Breeder | F(2, 130)=0.347, p=0.707 |
| | | | Poly(I:C) | F(1, 130)=1.081, p=0.300 |
| | | | Sex | F(1, 130)=0.215, p=0.644 |
| | | | Interaction(Breeder X Poly(I:C)) | F(2, 130)=0.214, p=0.807 |
| | | | Interaction(Breeder X Sex) | F(2, 130)=1.512, p=0.222 |
| | | | Interaction(Poly(I:C) X Sex) | F(1, 130)=2.885, p=0.092 |
| | | | Interaction<br>(Breeder X Poly(I:C) X Sex) | F(2, 130)=0.628, p=0.535 |
